# The TTLL10 polyglycylase is stimulated by tubulin glutamylation and inhibited by polyglycylation

**DOI:** 10.1101/2024.03.31.587457

**Authors:** Steven W. Cummings, Yan Li, Jeffrey O. Spector, Christopher Kim, Antonina Roll-Mecak

## Abstract

Microtubules in cells have complex and developmentally stereotyped posttranslational modifications that support diverse processes such as cell division, ciliary growth and axonal specification. Glycylation, the addition of glycines, singly (monoglycylation) or in chains (polyglycylation), is primarily found on axonemal microtubules where it functions in cilia maintenance and motility. It is catalyzed by three enzymes in the tubulin tyrosine ligase-like family, TTLL3, 8 and 10. We show that TTLL8 monoglycylates both α- and β-tubulin, unlike TTLL3 which prefers β-tubulin. Microscopy and mass spectrometry show that TTLL10 requires monoglycylation for high affinity microtubule binding and elongates polyglycine chains only from pre-existing glycine branches. Surprisingly, tubulin polyglycylation inhibits TTLL10 recruitment to microtubules proportional with the number of posttranslationally added glycines, suggesting an autonomous mechanism for polyglycine chain length control. In contrast, tubulin glutamylation, which developmentally precedes polyglycylation in cilia, increases TTLL10 recruitment to microtubules, suggesting a mechanism for sequential deposition of tubulin modifications on axonemes. Our work sheds light on how the tubulin code is written by establishing the substrate preference and regulation of TTLL glycylases and provides a minimal system for generating differentially glycylated microtubules for *in vitro* analyses of the tubulin code.

## INTRODUCTION

Microtubules are dynamic, non-covalent biopolymers essential for basic cellular processes in all eukaryotes. They serve as tracks for intracellular transport and build complex cellular structures such as the bipolar spindle and the axonemes in cilia and flagella. Microtubules are constructed of α/β-tubulin heterodimers which consist of a globular body, involved in polymerization interfaces, and intrinsically disordered C-terminal tails that decorate the microtubule outer surface, are heavily posttranslationally modified, and fulfill complex regulatory functions (1). Microtubule-based subcellular structures have distinct posttranslational modification patterns which regulate a microtubule’s interactions with effectors and its function in cellular physiology. This is known as the tubulin code (1, 2). Tubulin modification patterns are the result of the combinatorial action of tubulin modification enzymes which write and erase the tubulin code. Aberrant patterns of tubulin modifications are associated with tumorigenesis (3, 4), poor cancer prognosis (5), neurodegeneration (6, 7), neurodevelopmental disorders (8) and other pathologies (9), and mutations in tubulin modification enzymes are linked to several human pathologies (10). It is therefore crucial to understand the substrate specificity and regulation of the enzymes that introduce these modifications.

Ciliary microtubules are among the most heavily posttranslationally modified. Glycylation, which involves the addition of glycines, singly (monoglycylation) or in chains (polyglycylation) to internal glutamates in the C-terminal tails of α- or β-tubulin, appears largely limited to cilia where it is highly abundant (11) and where it accumulates as cilia mature (12, 13). Glycylation is ATP-dependent and catalyzed by three enzymes in the tubulin tyrosine ligase-like (TTLL) family, TTLL3, 8 and 10 (14, 15). Members of this family also catalyze glutamylation (16), which is also highly abundant in cilia and flagella. Glycylation is critical for the formation, stability, and function of cilia in protists (14, 17–19), zebrafish (14) and mammals (4, 12, 20, 21). Loss of the glycylase TTLL3 leads to a reduction in primary cilia and increased rates of cell division in colon epithelial cells (4), and TTLL3-inactivating mutations were identified in patients with colorectal cancer (4, 22). Loss of glycylation also severely impairs flagellar beating, resulting in male subfertility in mice (20). *In vitro* biophysical studies of microtubules isolated from Tetrahymena showed that glycylation increases microtubule stiffness (23). More recent work revealed that glycylation inhibits katanin microtubule severing (24). Thus, glycylation affects both the intrinsic biophysical properties of microtubules as well as how microtubules are recognized by effectors.

Glycylation is a polymodification. It involves two steps: initiation and elongation. During glycyl-initiation, a glycine is ligated to an internal glutamate residue through an isopeptide bond with the γ-carboxyl group. This is the branch-point glycine. In the initiation reaction, the internal glutamate residue is the acceptor amino acid, and the glycine residue is the donor amino acid (Figure 1A). In contrast, in the glycyl-elongation reaction, the glycine is ligated to the α-carboxyl group of an existing branch-point glycine or a polyglycine chain. In the elongation reaction, both the donor and the acceptor amino acids are glycines (Figure 1A). Thus, glycyl initiation and elongation necessitate the recognition of a glutamate acceptor and glycine acceptor, respectively, two amino acids with different physiochemical characteristics. Vertebrates have three glycylases: TTLL3, 8 and 10 (14, 15). Cellular overexpression studies coupled with the use of monoglycylation-specific antibodies (15) as well as biochemical and mass spectrometry-based studies (25) established that TTLL3 is a glycyl-initiase, adding only monoglycines to internal glutamates in tubulin tails, with a preference for β-tails (25). Cellular overexpression studies coupled with the use of antibodies that recognize mono- and polyglycylation indicate that TTLL8 is also a glycyl-initiase, while TTLL10 a glycyl-elongase (15, 26). However, direct biochemical evidence with purified enzymes for segregated initiation and elongation activity for glyclases is still lacking as does knowledge of their substrate specificity and regulation.

**Figure 1.**
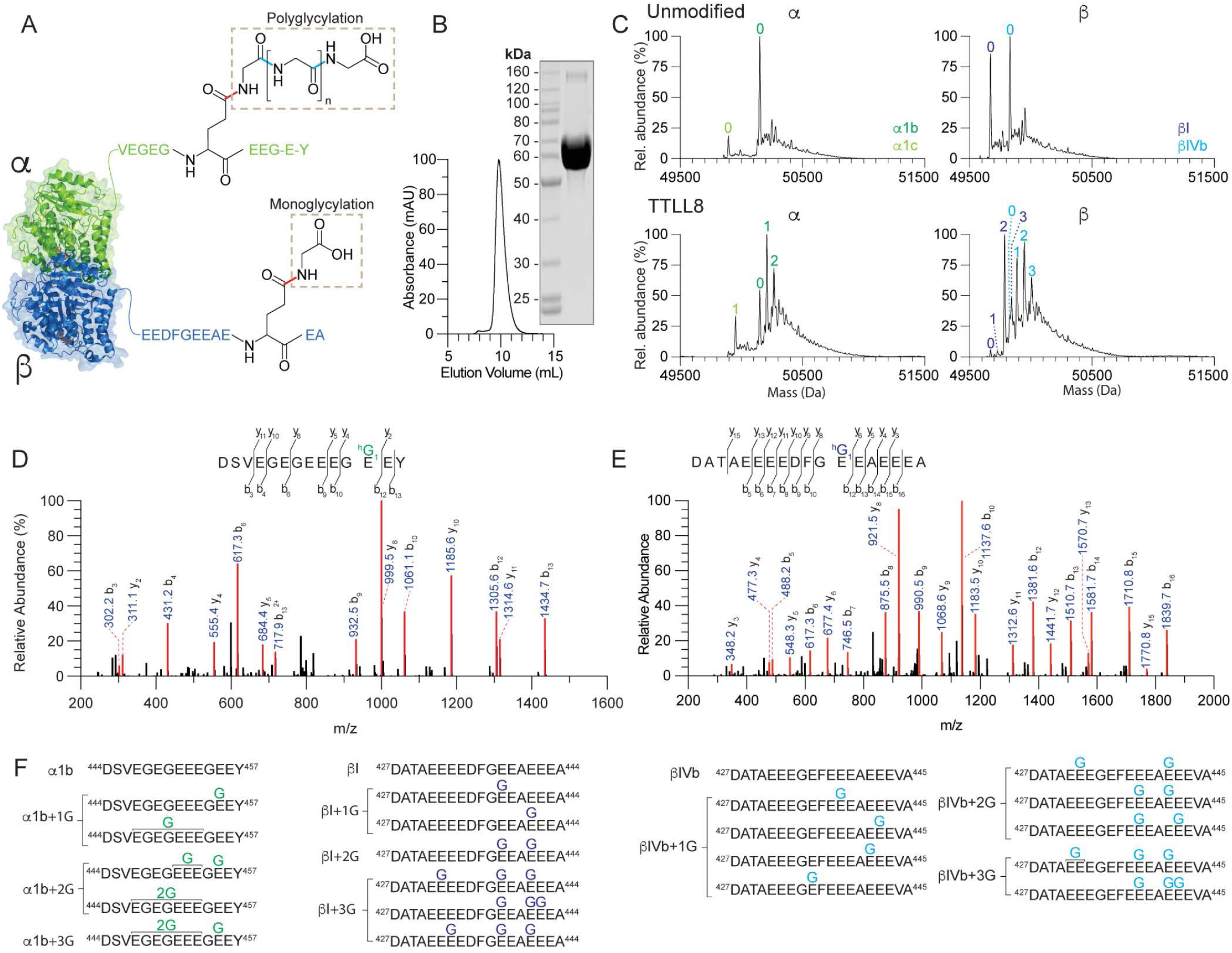
TTLL8 is a monoglycylase that initiates at multiple sites on α- and β-tubulin tails. (A) Cartoon of the αβ-tubulin dimer and C-terminal tail sequences illustrating the chemical structure of monoglycine branches and polyglycine chains. In monoglycylation, the donor (incoming) glycine is linked to the γ-carboxyl group of an acceptor glutamate in the tubulin tail *via* an isopeptide bond (red). In polyglycylation, the donor glycine is linked to the α-carboxyl group of the terminal acceptor glycine in the growing glycine chain *via* a standard peptide bond (cyan). (B) Size exclusion column elution profile and Coomassie-stained SDS-PAGE gel of recombinant purified TTLL8 (C) Mass spectra of unmodified human microtubules incubated without (top) or with TTLL8 (bottom). The number of posttranslationally added glycines is indicated in green (for α-tubulin isoforms) and blue (β-tubulin isoforms). The weighted mean of the number of glycines (<n^G^>) added to α- and β-tubulin are denoted α + <n^G^>; β + <n^G^>. For experimental details see Materials and Methods. (D-E) Representative MS/MS spectra from two independent experiments for monoglycylated α- (D) and β-tubulin (E) C-terminal tail peptide fragments proteolytically released from microtubules glycylated by TTLL8. For additional spectra see Figure S2. (F) List of TTLL8 monoglycylation sites identified by MS/MS sequencing. The line marked with 2G indicates that a monoglycine can be present at any of the glutamates in that region.

Using recombinant purified enzymes and well-defined tubulin substrates coupled with tandem mass spectrometry (MS/MS) we show that TTLL8 is exclusively a glycyl initiase, adding monoglycines to either α- or β-tubulin tails. Microscopy-based and MS/MS analyses show that TTLL10 requires monoglycylation for high-affinity microtubule binding and elongates polyglycine chains of variable lengths only from pre-existing monoglycines on both α- and β-tubulin tails. TTLL10 microtubule binding decreases monotonically with polyglycylation, indicating that TTLL10 activity is self-limiting and suggests an autonomous mechanism of polyglycine chain control. Furthermore, we find that TTLL10 microtubule recruitment is enhanced by glutamylation, a modification that precedes glycylation during cilia biogenesis (12), suggesting a potential biochemical basis for the order of deposition of these different tubulin modifications on the axoneme. Our work establishes the substrate preference and regulation of TTLL glycylases and sheds light on the biochemical mechanisms that generate tubulin modification patterns in cells.

## RESULTS AND DISCUSSION

### *In vitro* reconstitution of microtubule polyglycylation

To determine the substrate preference and biochemical activity of TTLL8, we purified recombinant human TTLL8 (Figure 1B) and assembled glycylation reactions with microtubules polymerized from unmodified human tubulin. Unmodified tubulin was isolated from tsA201 cells through a TOG affinity procedure. This tubulin consists primarily of unmodified α1b, βI and βIV-tubulin isoforms (27). Intact liquid chromatography mass spectrometry (LC-MS) of these reactions showed that TTLL8 adds glycines to unmodified microtubules on both α- and β-tubulin (Figure 1C). Using MS/MS we mapped glycylation sites on the TTLL8-modified microtubules (Figures 1D-F, S1, S2). The MS/MS analyses show that TTLL8 adds only branch-point glycines (monoglycines) to several glutamates on both the α- and β-tubulin tails (Figures 1D, 1E, 1F, S1, S2). A tubulin with more than one branch-point glycine is still referred as monoglycylated. We could not detect any polyglycine chains, indicating that TTLL8 is an initiase, also consistent with previous reports using antibody-based assays (15). TTLL8 activity was also detectable by Western blot which showed a saturated band after treatment of tubulin with TTLL8, consistent with the mass spectrometry data (Figure S3). Extracted ion chromatographs (XIC) for α1b peptides show a dominant peak at position E455 (Figures S1A) with additional possible modifications at positions E447-E453 (Figures 1F, S1A-C, S2A). XIC for βI peptides show a dominant peak at position E438 followed by a less abundant peak at position E441 (Figure S1D). Monoglycylation at these two sites yields the most abundant species with two monoglycines (Figure S1E). Interestingly, modification at E432 is found in the most abundant species with three or four monoglycines (Figures S1F, S1G, S2B), however modification at this position is not abundant in either the +1 or the +2 glycine peaks, indicating the lower preference of the enzyme for this site, or the possible dependence on modification at other sites prior to activity at the E432 site. Similar modification patterns were observed on βIVb (Figure S1H). Thus, TTLL8 is an initiase that adds monoglycines at multiple positions in both α- and β-tubulin tails. Our previous work with recombinant tubulin TTLL3 showed that enzyme to also be strictly a monoglycylase (25).

We then purified recombinant *Xenopus tropicalis* TTLL10 (Figure 2A) and used it in *in vitro* assays with unmodified microtubules. In contrast to TTLL8, TTLL10 glycylates only microtubules that are already monglycylated and does not glycylate unmodified microtubules, even when incubated for a prolonged time (Figure 2B). This is consistent with cellular overexpression data which showed that polyglycylation signal was detected *via* antibody only in tubulin from cells that co-expressed TTLL8 and TTLL10, but not TTLL10 alone (15, 26). LC-MS analyses show that TTLL10 can add glycines to both α- and β-tubulin as long as they were previously monoglycylated (Figure 2B). MS/MS shows that TTLL10 is only elongating glycine chains from pre-existing monoglycines and does not initiate new glycine chains (Figures 2C-E, S4A-C, S5A-B). To facilitate the MS/MS analyses we used heavy glycine in our modification reactions to distinguish between the TTLL10 added glycines and pre-existing glycines on tubulin (Materials and Methods). XIC analysis indicates that, consistent with βI position E438 and α1B position E455 being the dominant sites modified by TTLL8, the main species in TTLL10 modified microtubules contain polyglycine chains of variable lengths at α1B position E455 and βI position E438 (Figures S4A, S4B, S5A-B). Thus, TTLL10 elongates polyglycine chains from pre-existing monoglycines at multiple positions on α and β-tubulin tails.

**Figure 2.**
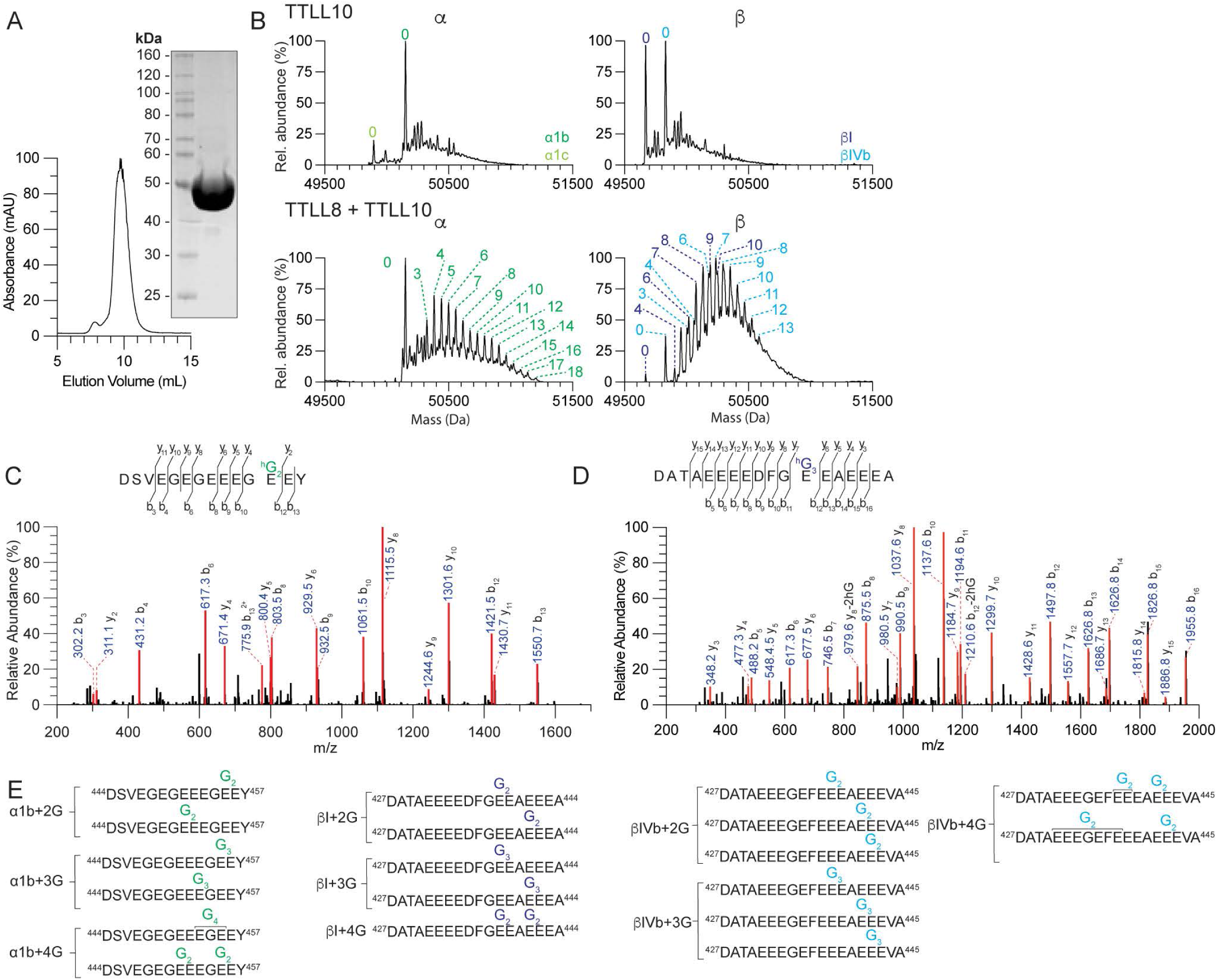
TTLL10 is exclusively a tubulin polyglycylase that can only elongate existing branch-point glycines. (A) Size exclusion column elution profile and Coomassie-stained SDS-PAGE gel of recombinant purified TTLL10. (B) Deconvoluted α- and β-tubulin mass spectra from unmodified human microtubules incubated with TTLL10 alone or with TTLL8 followed by incubation with TTLL10. The weighted mean of the number of glycines is denoted as in Figure 1. (C-D) Representative MS/MS spectra from two independent experiments for polyglycylated α- (C) and β-tubulin isoforms (D) C-terminal tail peptide fragments proteolytically released from microtubules glycylated by TTLL8 and TTLL10. For additional spectra see Figure S5. (E) List of TTLL10 polyglycylation sites identified by MS/MS sequencing.

### TTLL10 recognizes monoglycines on tubulin tails

Next, we wanted to understand the mechanistic basis for the exclusive activity of TTLL10 on microtubules that already had monoglycine branches. We used total internal reflection fluorescence (TIRF) microscopy to visualize the association of Alexa647-labeled TTLL10-SNAP with unmodified and TTLL8-modified microtubules (Materials and Methods; Figure 3A and Figure S6). These assays show that TTLL10 binds robustly to monoglycylated microtubules (Figure 3B), with an apparent K_d_ of ∼ 600 nM, while showing little binding to unmodified microtubules. Moreover, assays with microtubules with different monoglycylation levels revealed that TTLL10 recruitment to the microtubule increases with increasing numbers of monoglycines on tubulin (Figures 3C, D). Specifically, TTLL10 recruitment to monoglycylated microtubules (<n^G^>_α_ ∼ 0.8, <n^G^>_β_ ∼ 1.2) increases more than ∼ sixfold compared to unmodified microtubules. Addition of more monoglycines to the tubulin tails has a more muted effect, resulting in only a ∼17% increase in recruitment to microtubules with <n^G^>_α_ ∼ 1.1, <n^G^>_β_ ∼ 2.5 (Figure 3D). This suggests that the TTLL10 active site recognizes a single branch-point glycine and that steric hindrance on the tubulin tails limits the number of branch-point glycines that can be simultaneously recognized by multiple TTLL10 enzymes. The strong stimulation of TTLL10 recruitment to the microtubule by monoglycylation is consistent with our activity assays that show TTLL10 adds glycines only to microtubules that already have branch-point glycines on their C-terminal tails (Figure 2). This mechanism of branch-point glycine recognition contrasts with that of the glutamyl elongase TTLL6 which binds to unmodified and glutamylated microtubules with similar affinity, but has a higher catalytic rate on tubulin tails that have pre-existing branch-point glutamtates (28). The two-enzyme requirement for the addition of polyglycine chains to unmodified tubulin is also consistent with the chemically distinct substrates that mono- and polyglycylation requires: addition to a glutamate first (monoglycylation), followed by addition to a glycine (polyglycylation; Figure 1A) in contrast to glutamylation which uses a glutamate as a substrate for both the initiation and elongation steps.

**Figure 3:**
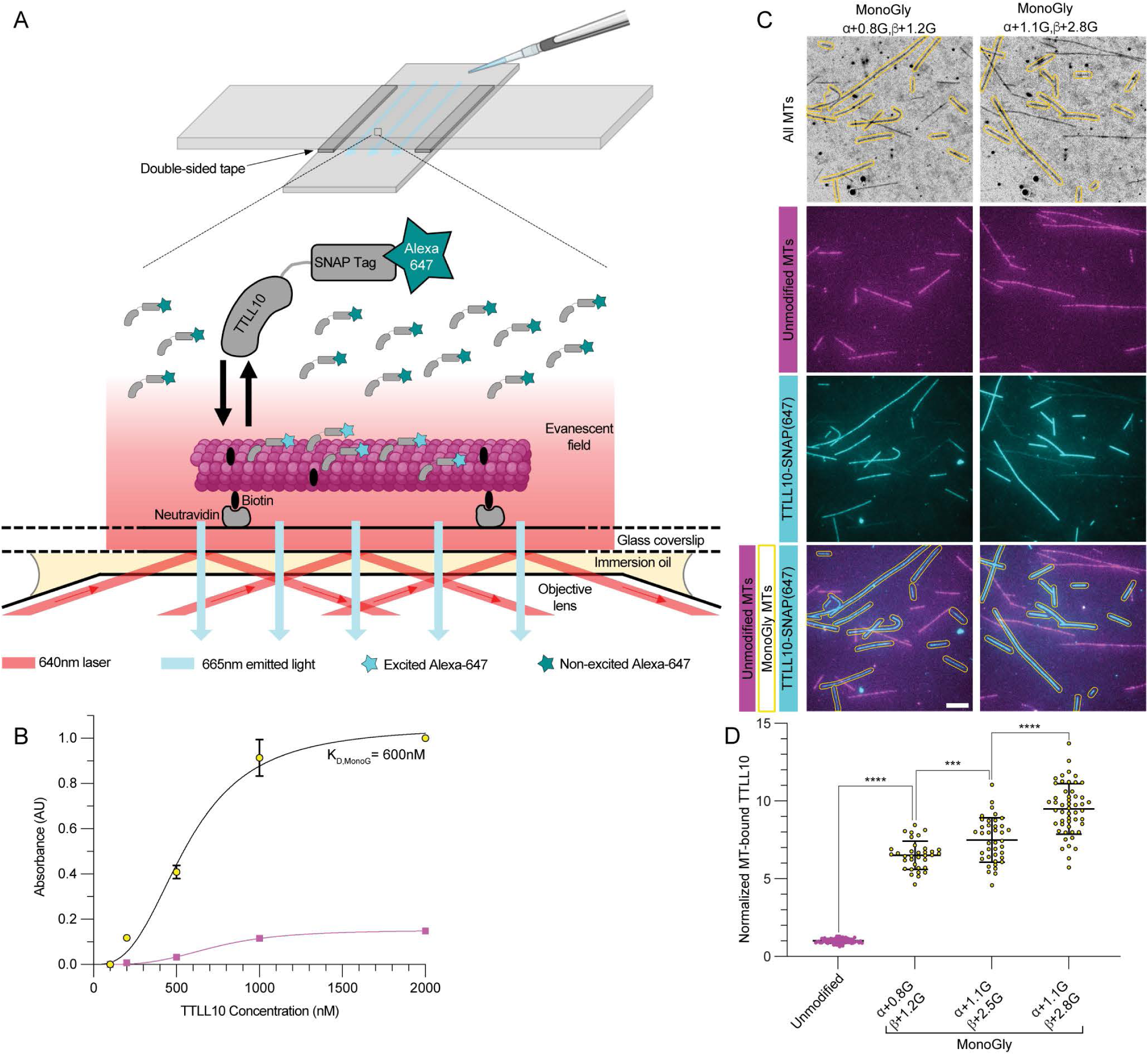
TTLL10 requires monoglycylation for robust microtubule binding. (A) Diagram of TIRF-based microtubule binding assays (B) Binding curve of TTLL10-SNAP(Alexa647) to unmodified microtubules, shown in magenta, and monoglycylated microtubules (α + 1.7G, β + 2.0G), shown in yellow with black outline. It was not possible to reach saturation in the reaction with unmodified microtubules. (C) TTLL10-SNAP(Alexa647) (cyan) association with unmodified microtubules (Hi-Lyte 488-labeled, magenta) and monoglycylated microtubules (unlabeled, yellow contour). The weighted mean of the number of glycines denoted as in Figure 1. For LC-MS of microtubules used in this assay see Figure S6. Assays performed at 500 nM TTLL10. Scale bar, 5 μm. (D) Quantification of TTLL10-SNAP(647) recruitment to differentially monoglycylated microtubules. Values normalized to unmodified microtubules; n = 66, 35, 39, and 51 microtubules from 14, 3, 5, and 6 independent experiments for the unmodified, β+1.2G, β+2.5G, and β+2.8G conditions, respectively. Statistical significance determined by Welsh’s t-test, with p < 0.0001 (****) or p < 0.001 (***).

### TTLL10 activity on microtubules is self-limiting

Given the pronounced effect monoglycylation has on TTLL10 recruitment to microtubules, we then investigated the effect of polygylcylation on TTLL10 microtubule binding. For this, we generated microtubules with different polyglycylation levels (Figure S7) and used them in TIRF-based assays. These showed that TTLL10 recruitment to polyglycylated microtubules is dramatically reduced compared to monoglycylated microtubules (Figures 4A, 4B). Strikingly, the binding decreases monotonically with polyglycine chain length (Figure 4B). This contrasts with monoglycylation which is stimulatory for TTLL10 recruitment (Figures 3B, 3C, 3D). We observed a similar inhibition of TTLL10 microtubule recruitment by polygycylation when we monitored TTLL10 in the microscopy chambers as the glycylation reaction was ongoing in the presence of ATP and glycine. These experiments showed that TTLL10 recruitment to monoglycylated microtubules quickly peaks after addition of TTLL10 into the chamber and then decreases, as the polyglycylation reaction proceeds, to a level close to that observed for unmodified microtubules (Figures 4C, 4D). In contrast, when this assay is performed in the absence of glycine, precluding the elongation of polyglycine chains, TTLL10 binding to monoglycylated microtubules plateaus and does not decrease with time (Figure 4D). TTLL10 binding to unmodified microtubules was close to background regardless of whether free glycine was present or absent. These results indicate that the gradual loss of TTLL10 from the microtubule is caused by its polyglycylation activity, consistent with our results examining TTLL10 binding to differentially polyglycylated microtubules (Figures 4A, 4B). Polyglycine as well as polyglutamate chain length regulate the activity of effectors such as kinesin (29) and microtubule severing enzymes (24, 30). Therefore, chain length regulation is important for controlling cellular responses. While glutamylation levels are maintained by the balance between TTLL glutamylases, which add glutamate chains (31), and CCP carboxypeptidases which remove them (32–36), an enzyme that processes glycine chains or branches has yet to be identified, raising the question of how polyglycine chain lengths are regulated in cells. The inhibition of TTLL10 binding by tubulin polyglycylation that we uncovered suggests a possible mechanism for limiting polyglycine chains lengths on tubulin, independent of a deglycylating enzyme.

**Figure 4:**
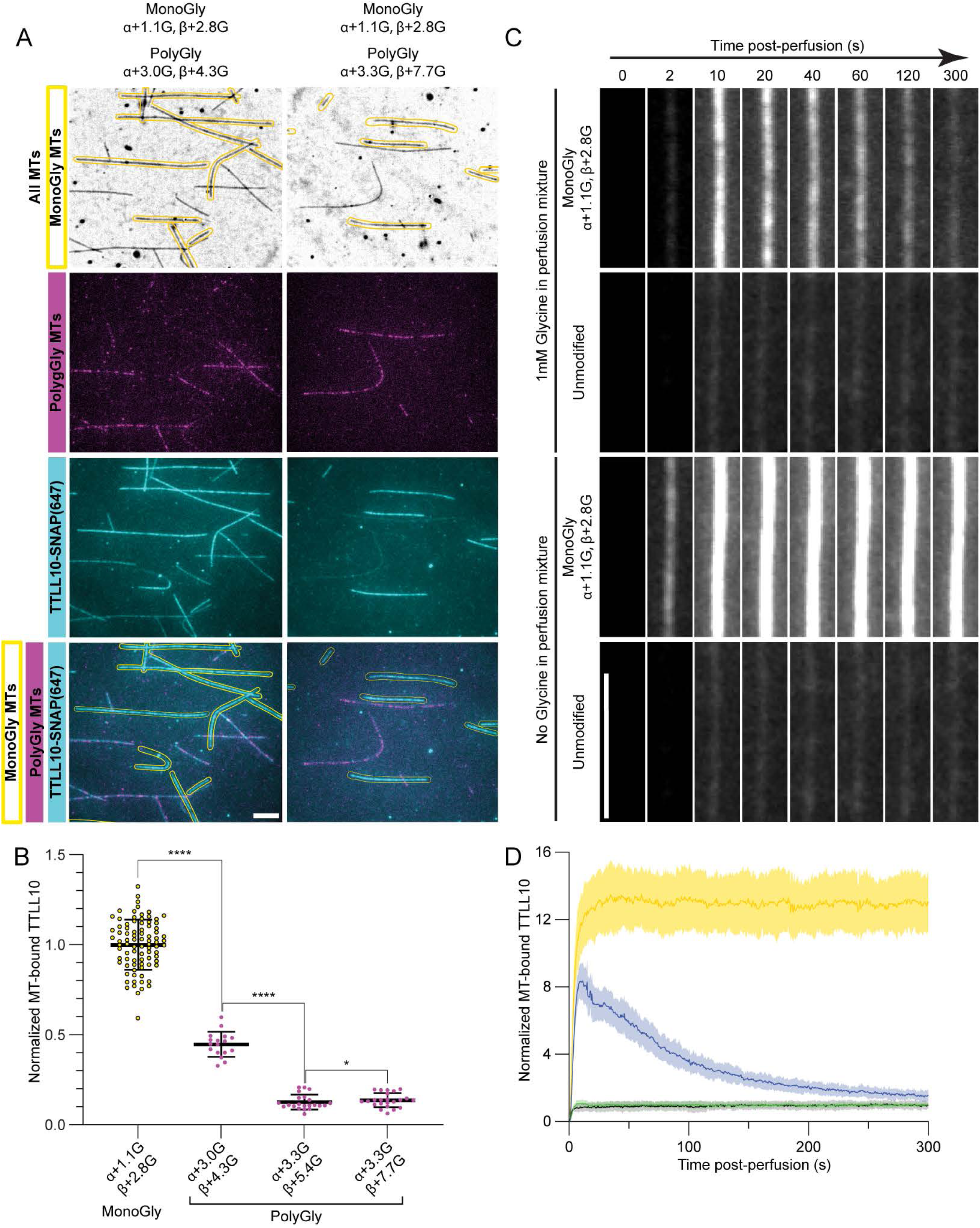
TTLL10 recruitment to microtubules decreases with increased polyglycylation. (A) TTLL10-SNAP(647) (cyan) association with monoglycylated microtubules (unlabeled, outlined in yellow) and polygylcylated microtubules (HiLyte 488-labeled, magenta). The weighted mean of the number of glycines denoted as in Figure 1. For LC-MS of microtubules used in this assay see Figure S7. Assays performed at 500 nM TTLL10. Scale bar, 5 μm. (B). Quantification of TTLL10-SNAP(Alexa647) recruitment of differentially polyglycylated microtubules normalized to levels on monoglycylated microtubules; n = 59, 20, 28, and 33 microtubules from 10, 3, 3, and 4 independent experiments for the β+2.8G, β+4.3G, β+5.4G, and β+7.7G conditions, respectively. Statistical significance determined by Welsh’s t-test, with p < 0.0001 (****) or p < 0.05 (*). (C) Representative images showing TTLL10-SNAP(647) association with monoglycylated and unmodified human microtubules as a function of time, in the presence of 1mM ATP with 1mM glycine (top two panels) and without 1mM glycine (bottom two panels). Scale bar, 5 μm. (D) Time courses of TTLL10-SNAP(Alexa647) recruitment to microtubules monoglycylated by TTLL8 (<n^G^>α ∼ 1.1, <n^G^>β ∼ 2.8) in the presence of 1 mM ATP with 1mM glycine (blue), 1 mM ATP without 1mM glycine (yellow), and unmodified microtubules in the presence of 1 mM ATP with 1mM glycine (black), and without 1mM glycine (green); n = 12 monoglycylated and 7 unmodified microtubules for the 1mM glycine time courses, and 8 monoglycylated and 8 unmodified microtubules for the glycine-free time courses. The same effect was observed across 3 independent experiments.

### Glutamylation increases TTLL10 recruitment to microtubules

Axonemal microtubules are abundantly glutamylated. Glutamylation appears during cilia development first, followed by glycylation (12, 13), indicating that in this scenario glycylases act on pre-glutamylated microtubule substrates. TTLL6, an α-tubulin specific glutamyl elongase (28, 31, 37) localizes to cilia where it is required for their biogenesis and for normal beat patterns (12, 38). We therefore sought to determine the effect of glutamylation by TTLL6 on TTLL10 microtubule binding. For this we generated microtubules that were either only monoglycylated or both monoglycylated and polyglutamylated. Our mass spectra as well as Western blots indicate that the tubulin purified from tSA201 cells has no detectable glutamylation (Figures 1A, S8). To ensure the exact same monoglycylation levels in our assays, we first monoglycylated the microtubules. Then, we divided the sample in two: one half was glutamylated by incubation with TTLL6, while the other half was incubated with TTLL6 in the absence of glutamate (Materials and Methods). Thus, both samples had the exact same monoglycylation levels (<n^G^>_α_ ∼ 0.1, <n^G^>_β_ ∼ 0.8) and differed only in their glutamylation status. TIRF-based assays with these substrates (Figure 5A) showed that polyglutamylation by TTLL6 (<n^E^>_α_ ∼ 18, <n^E^>_β_ ∼ 4) results in a ∼1.8-fold increase in TTLL10 recruitment to monoglycylated microtubules (Figure 5B, Figure S9). Glutamate chains as long as ∼ 17 glutamates have been documented on α-tubulin from axonemes (39). The stimulation of microtubule binding by polyglutamylation also applied to polyglycylated microtubules: TTLL10 binding to polyglycylated microtubules that were also glutamylated was ∼3.2-fold higher than to microtubules that were only polyglycylated (Figure 5B). Thus, glutamylation enhances the recruitment of TTLL10 to microtubules. Polyglycylation appears later in cilia development, after polyglutamylation. Given the moderate binding affinity of TTLL10 for monoglycylated microtubules (apparent k_d_ ∼ 600 nM) and the low expression levels of TTLL10 (12), we speculate that TTLL10 is not recruited effectively to microtubules until they have been glutamylated first, ensuring the sequential addition of these modifications during cilia maturation.

**Figure 5:**
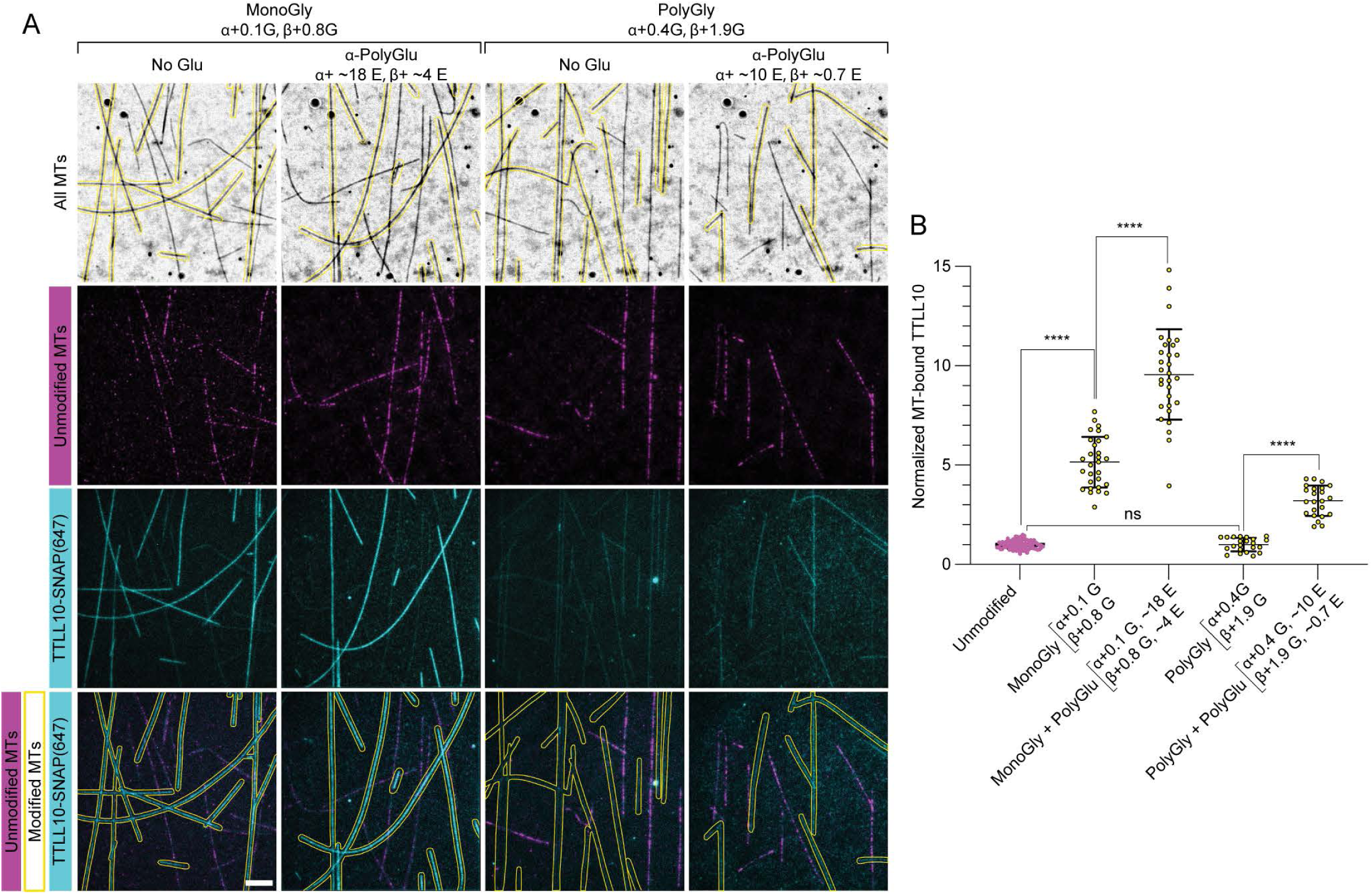
Polyglutamylation by TTLL6 enhances TTLL10 microtubule association. (A) TTLL10-SNAP(Alexa647) (cyan) association with unmodified microtubules (HiLyte 488-labeled, magenta) and differentially modified microtubules (unlabeled, outlined in yellow): monoglycylated, monoglycylated/polyglutamylated, polyglycylated only, and polygylcylated/polyglutamylated. Monoglycylation added by TTLL8, polyglycylation by TTLL10 and polyglutamylation by TTLL6 (Materials and Methods). The weighted mean of the number of glycines denoted as in Figure 1. Glutamylation levels for glycylated/glutamylated microtubules are approximate and estimated by western blot (Figure S9) because quantification from LC-MS spectra of these dual-modified microtubules was not possible due to the large number of species present. Scale bar, 5 μm. (B) Quantification of TTLL10-SNAP(Alexa647) recruitment to modified microtubules normalized to those on unmodified microtubules; n = 84, 29, 30, 22, and 23 microtubules from 10, 4, 4, 3, and 3 independent experiments for monoglycylated, monoglycylated/polyglutamylated, polyglycylated, and polyglycylated/polyglutamylated microtubules, respectively. Statistical significance determined by Welsh’s t-test, with p < 0.0001 (****).

In conclusion, using *in vitro* reconstitution we demonstrate that TTLL8 is strictly a glycyl initiase that catalyzes the addition of monoglycines at multiple positions on both α and β-tubulin tails, while TTLL10 is exclusively a tubulin glycyl elongase that catalyzes the addition of glycines only to pre-existing glycine branches on either α or β-tubulin. We note that TTLL glycylases seem to be more promiscuous in their substrate recognition, unlike TTLL glutamylases which show stronger preference for α or β-tubulin tails (28, 31, 40, 41). This difference in substrate stringency is consistent with the larger diversification of TTLL glutamylases (nine total identified in vertebrates so far), compared to glycylases. TTLL10 binding to microtubules requires priming of the tubulin tail with monoglycylation, establishing a hierarchy of enzyme recruitment to the microtubule. TTLL10 binding is progressively inhibited by polyglycylation, proportional with polyglycine chain length, suggesting an autonomous mechanism of polyglycine chain length control. TTLL10 recruitment to microtubules is stimulated by TTLL6 polyglutamylation, demonstrating the interplay between these two modifications. Because polyglycylation inhibits the association of TTLL10 with the microtubule, the stimulatory effect of glutamylation on TTLL10 recruitment can potentially generate longer polyglycyine chains on microtubules that are already glutamylated and shorter polyglycine chains on microtubules that are not glutamylated. The stimulatory effect of glutamylation does not necessarily result in increased glycylation of the same tubulin dimer that is glutamylated because TTLLs can interact with the tubulin tails of neighboring tubulin dimers on the microtubule (40), and also because increased recruitment to one glutamylated tubulin subunit ensures a higher probability of rebinding at a nearby tubulin on the microtubule. These mechanisms can result in an enrichment of glycylation on the same microtubule, but not necessarily on the same tubulin dimer that carries the stimulatory polyglutamate chains on its tail. Recent work revealed that glutamylation also stimulates vasohibin (42), offering a possible explanation for the high degree of overlap between detyrosination and glutamylation on subpopulations of microtubules in cells. More recently, the α-tubulin glutamylase TTLL6 was shown to read out the modification status of the β-tubulin tail on the laterally adjacent neighboring tubulin dimer, facilitating crosstalk between tubulin tails (43). Our present study adds to the growing evidence of crosstalk between tubulin modifications and establishes the biochemical mechanisms that govern the syntax of the tubulin code. Lastly, a major limitation towards our mechanistic understanding of glycylation and the tubulin code overall has been the inability to generate microtubules with diverse and controlled posttranslational modifications patterns for *in vitro* reconstitution. By identifying the sites of modifications introduced by TTLLs glycylases and the interplay between their activities, we not only provide insight into the biochemical program that generates the complex *in vivo* microtubule glycylation patterns, but also a tool for generating differentially glycylated microtubules to study *in vitro* the effects of glycylation on microtubule effectors and thus shed light on the mechanism of action of the tubulin code.

## Acknowledgements

We thank Duck-Yeon Lee at the National Heart, Lung, and Blood Institute (NHLBI) Biochemistry Core for mass spectrometer access. This research was supported by the Intramural Research Program of the National Institutes of Health (NIH). The contributions of the NIH author(s) were made as part of their official duties as NIH federal employees, are in compliance with agency policy requirements, and are considered Works of the United States Government. However, the findings and conclusions presented in this paper are those of the author(s) and do not necessarily reflect the views of the NIH or the U.S. Department of Health and Human Services.

## Author contributions

SWC expressed and purified proteins, did LC-MS and microscopy assays; YL did MS/MS, JOS did microscopy assays together with SWC. SWC and JOS did image analysis. CK did initial expression and purification of TTLL8. SWC, YL, JOS analyzed data. SWC and ARM planned experiments, discussed data and wrote the manuscript. ARM conceived the project and supervised research. All authors reviewed and approved the manuscript.

## Data availability

All plasmids used in this study are available from the corresponding author upon request and source data will be provided for each figure.

---

**Figure S1.**
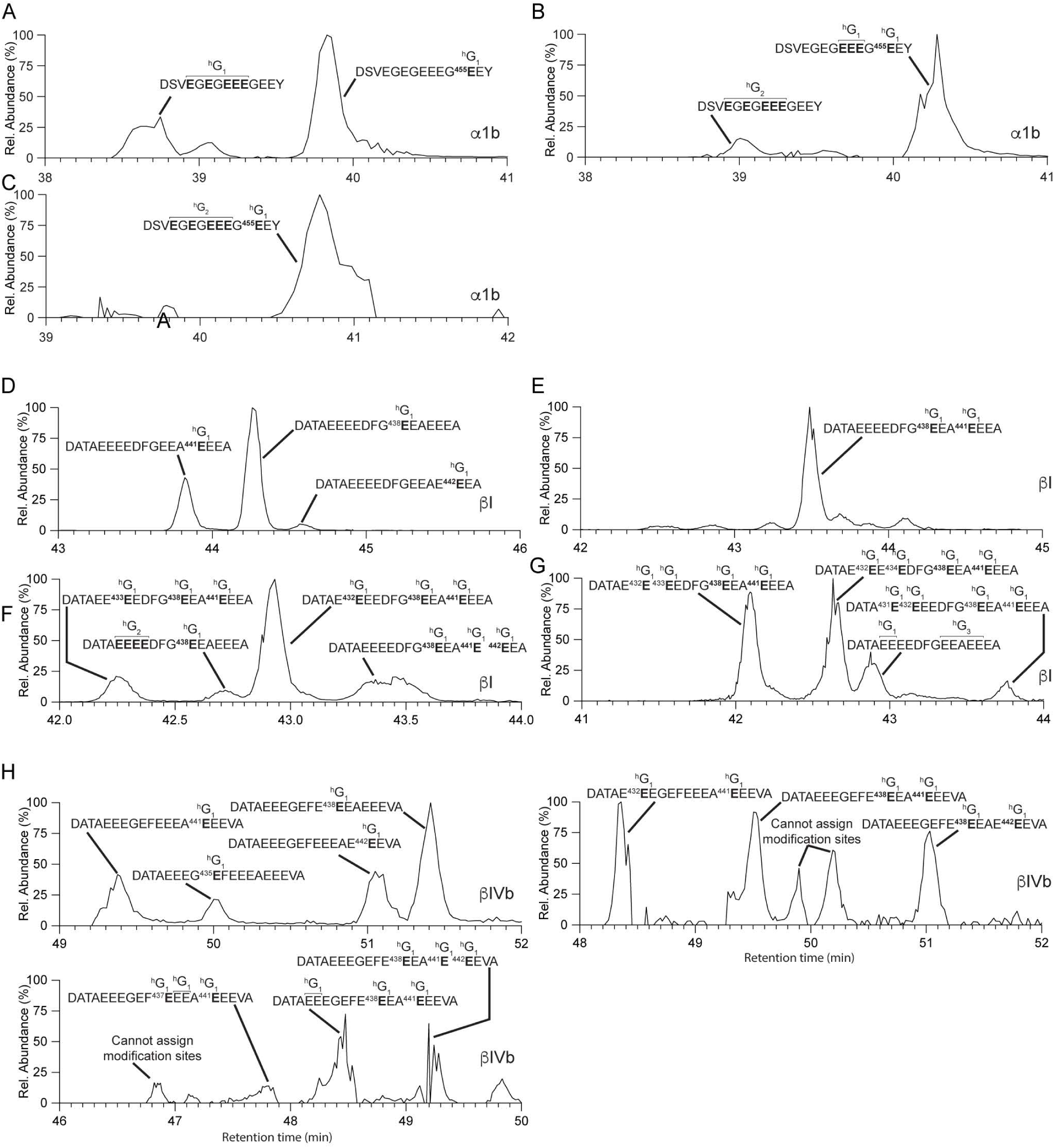
Extracted-ion chromatograms for monoglycylated α1b, βI, and βIVb tubulin tails modified by TTLL8. (A-C). Extracted-ion chromatograms for monoglycylated α1b-tail peptides with one monoglycine branch (A), two monoglycine branches (B) and three monoglycine branches (C). (B-G). Extracted-ion chromatograms for monoglycylated βI-tail peptides with one monoglycine branch (D), two monoglycine branches (E), three monoglycine branches (F) and four monoglycine branches (G). (H). Extracted-ion chromatograms for monoglycylated βIVb-tail peptides. Tubulin tail peptides were proteolytically released from microtubules incubated with TTLL8.

**Figure S2.**
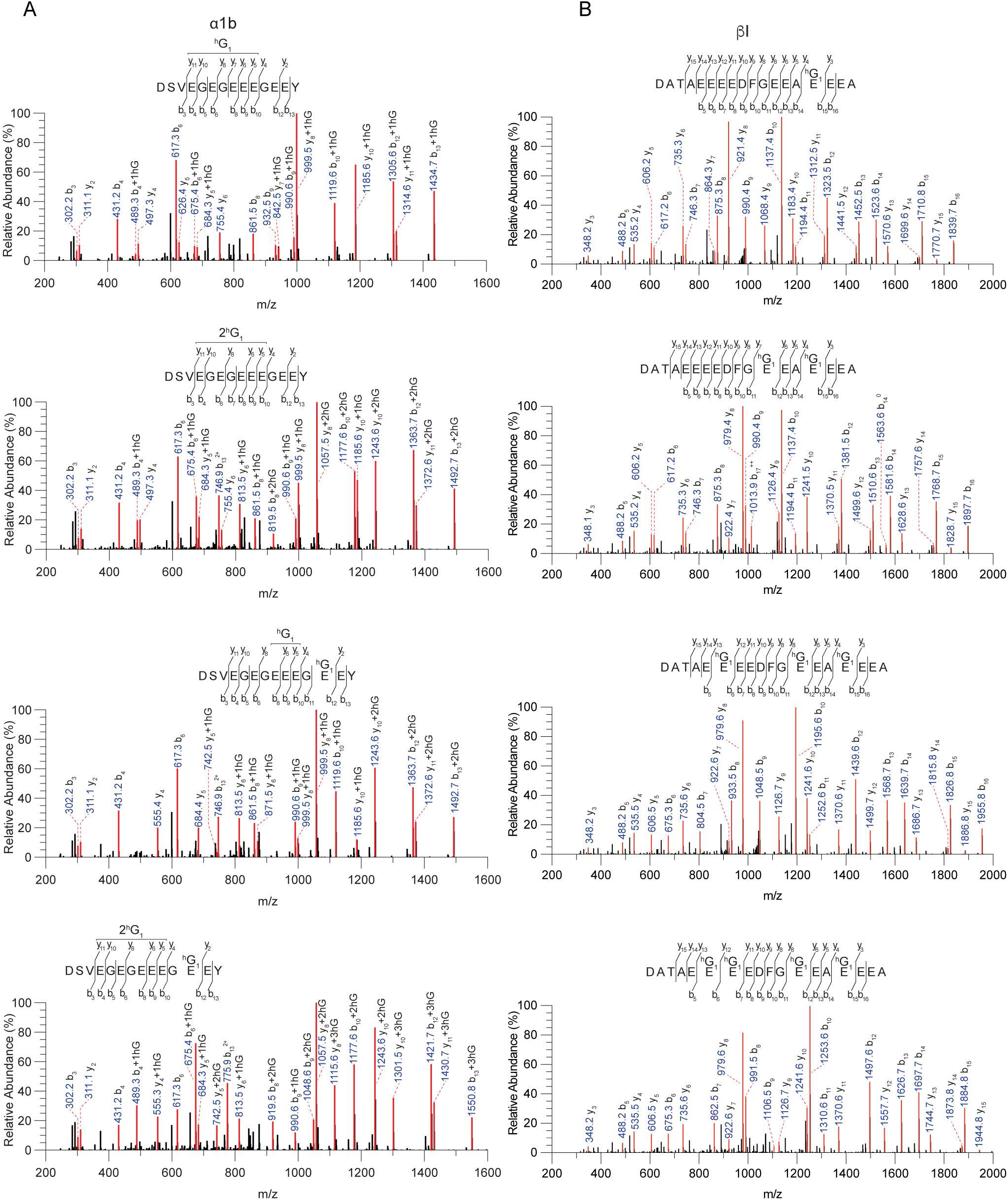
MS/MS spectra for monoglycylated tubulin C-terminal tail peptides proteolytically excised from microtubules incubated with TTLL8. (A, B). MS/MS spectra for monoglycylated α1b-peptides (A) and βI-tubulin C-terminal tail peptides (B).

**Figure S3.**
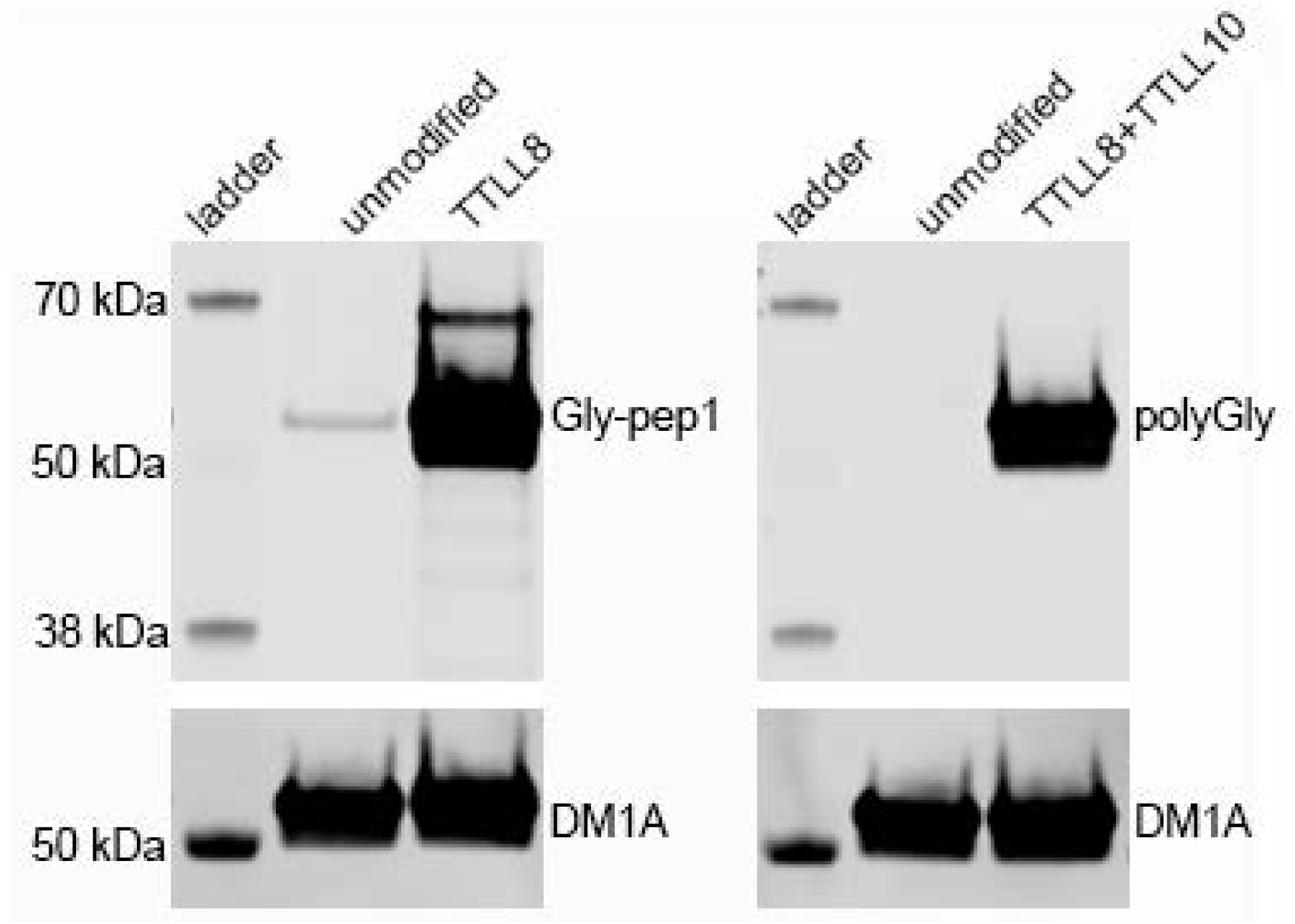
Western blot analysis of glycylation in tubulin purified from tSA201. 0.2 μgs of purified tubulin were loaded for each unmodified and glycylated sample. Glycylation was detected with the Gly-pep1 and anti-polyG antibodies (Materials and Methods). We note a weak reactivity to the gly-pep1 antibody for the tubulin purified from tSA201 cells. It is unclear whether the antibody detects a very small proportion of mono- or bi-glycylated tubulin in this sample or the signal is due to a low affinity interaction of the antibody for the unmodified tubulin tail which is visible at these tubulin loading levels. Neither our LC-MS or MS/MS data of the tsA201 tubulin detected any glycylation on the intact tubulin, or tubulin tails, respectively. The higher molecular weight band is from TTLL8 which self-modifies during expression. The tSA201 tubulin shows no reactivity against the polyglycylation antibody.

**Figure S4.**
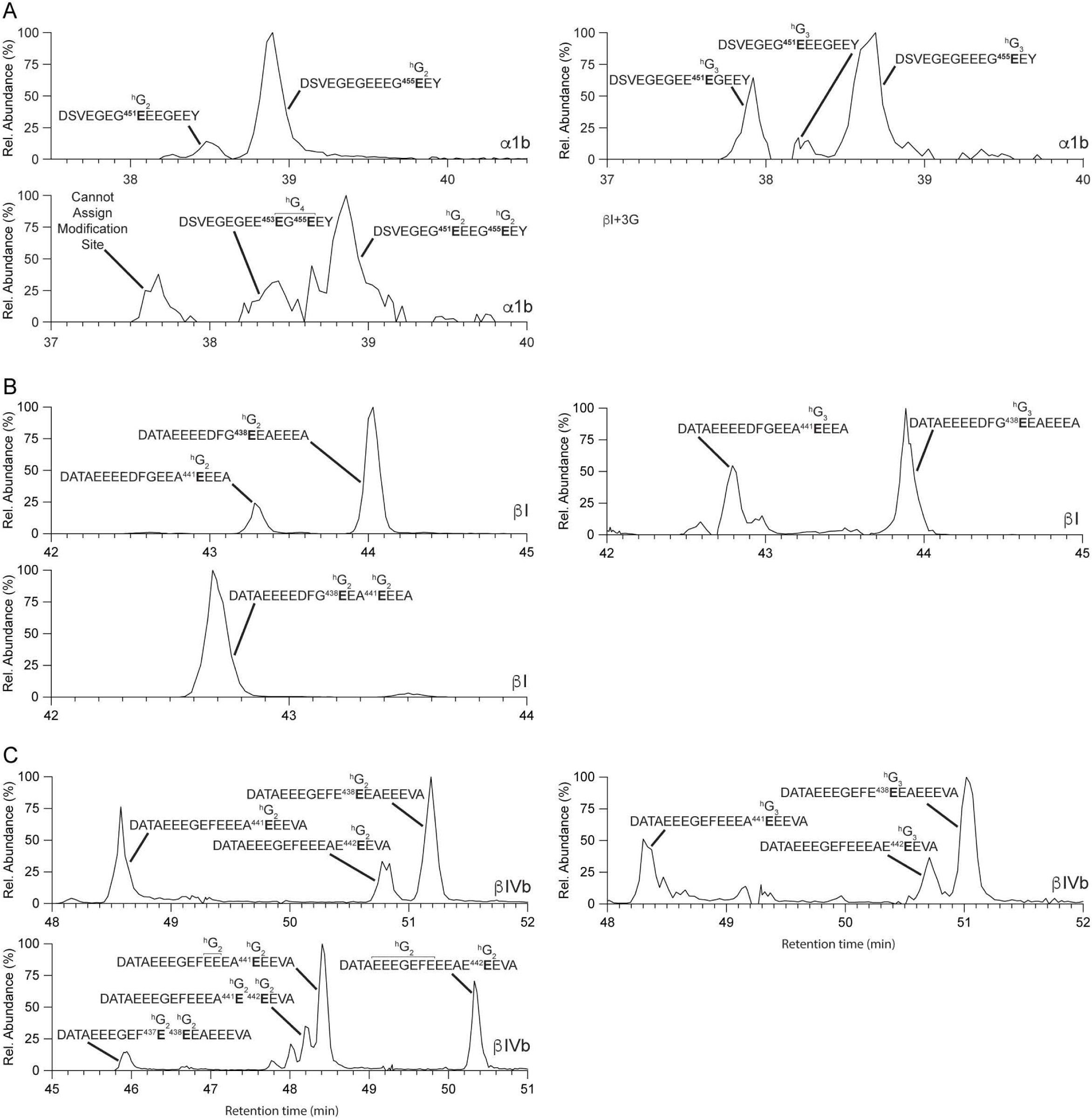
Extracted-ion chromatograms for polyglycylated tubulin C-terminal tail peptides proteolytically excised from monoglycylated microtubules incubated with TTLL10. (A-C). Extracted-ion chromatograms for polyglycylated α1b-peptides (A), βI-peptides (B) and βIVb-tubulin (C) C-terminal tail peptides. The subscript in G_i_ indicates the length of the polyglycine chain.

**Figure S5.**
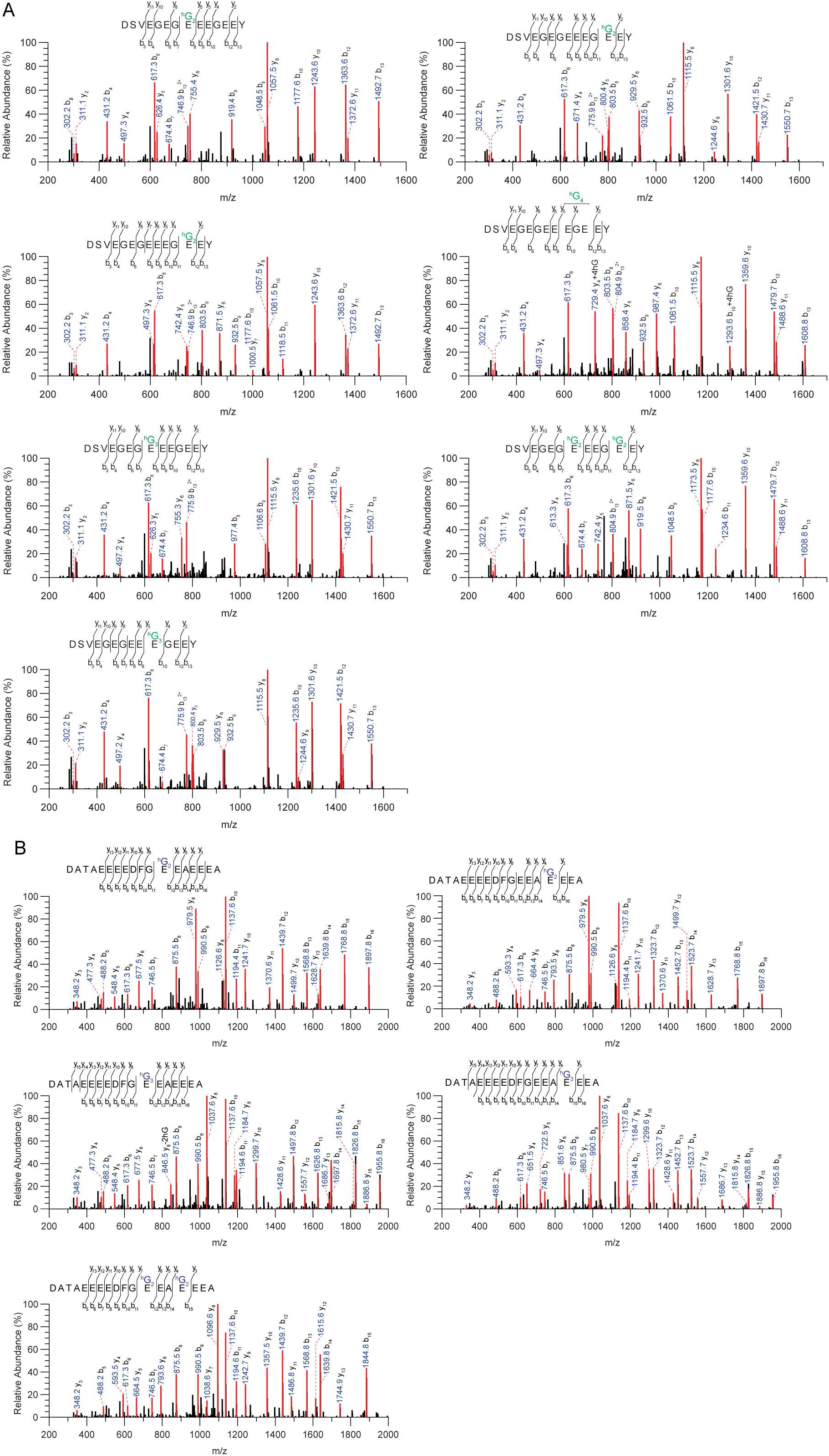
MS/MS spectra for polyglycylated tubulin C-terminal tail peptides proteolytically released from microtubules incubated with TTLL10 show elongation of polyglycine chains only at positions where monoglycine branches were already initiated by TTLL8. (A, B). MS/MS spectra for α1b-(A) and βI-tubulin (B) C-terminal tail peptides. The subscript in G_i_ indicates the length of the polyglycine chain.

**Figure S6.**
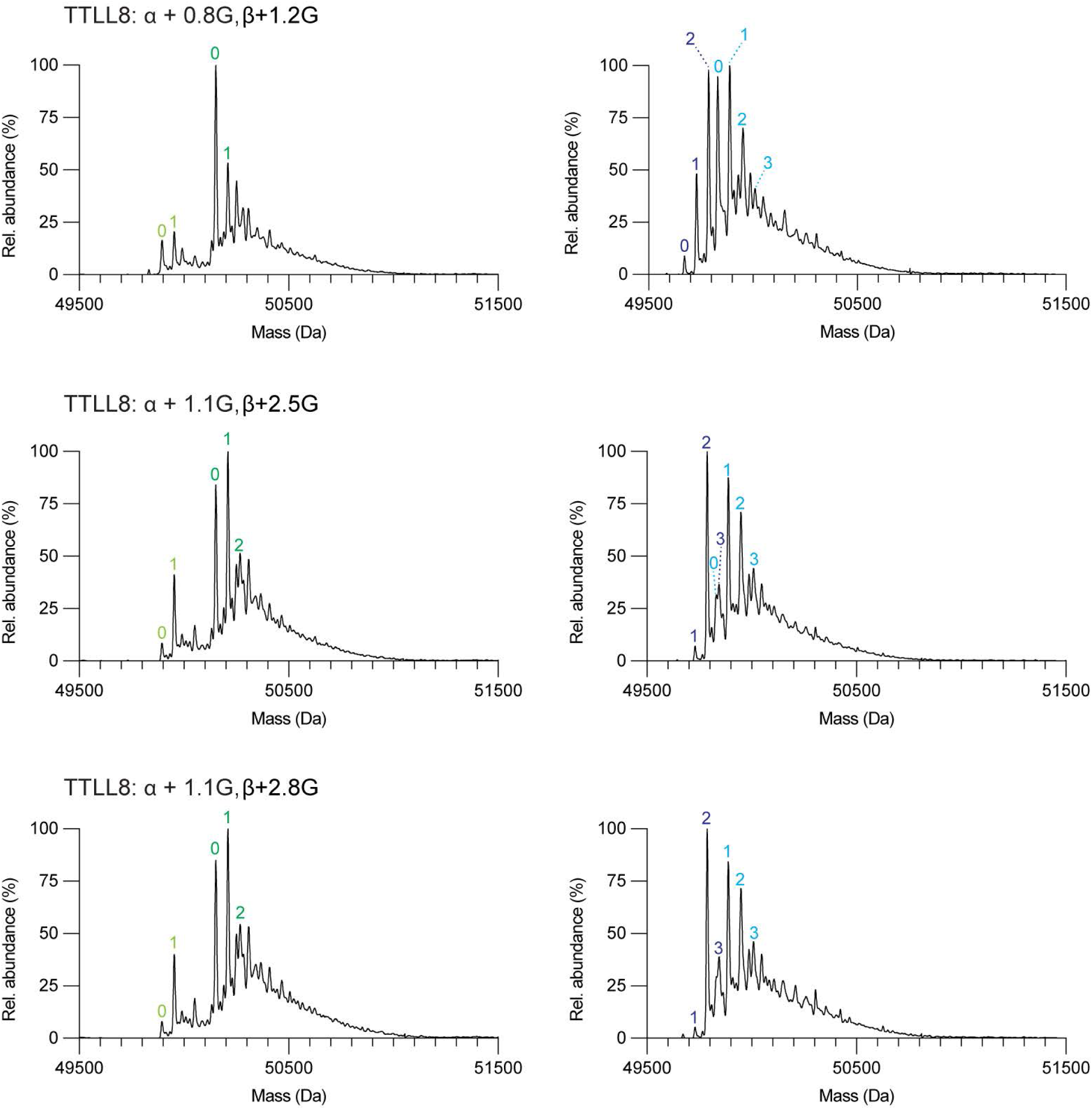
LC-MS of differentially monoglycylated microtubules used in assays shown in Figure 3. The weighted mean of the number of glycines (<n^G^>) added to α- and β-tubulin are denoted α + <n^G^>; β + <n^G^>. The number of posttranslationally added glycines is indicated in green (for α-tubulin isoforms) and blue (β-tubulin isoforms) on the spectra.

**Figure S7.**
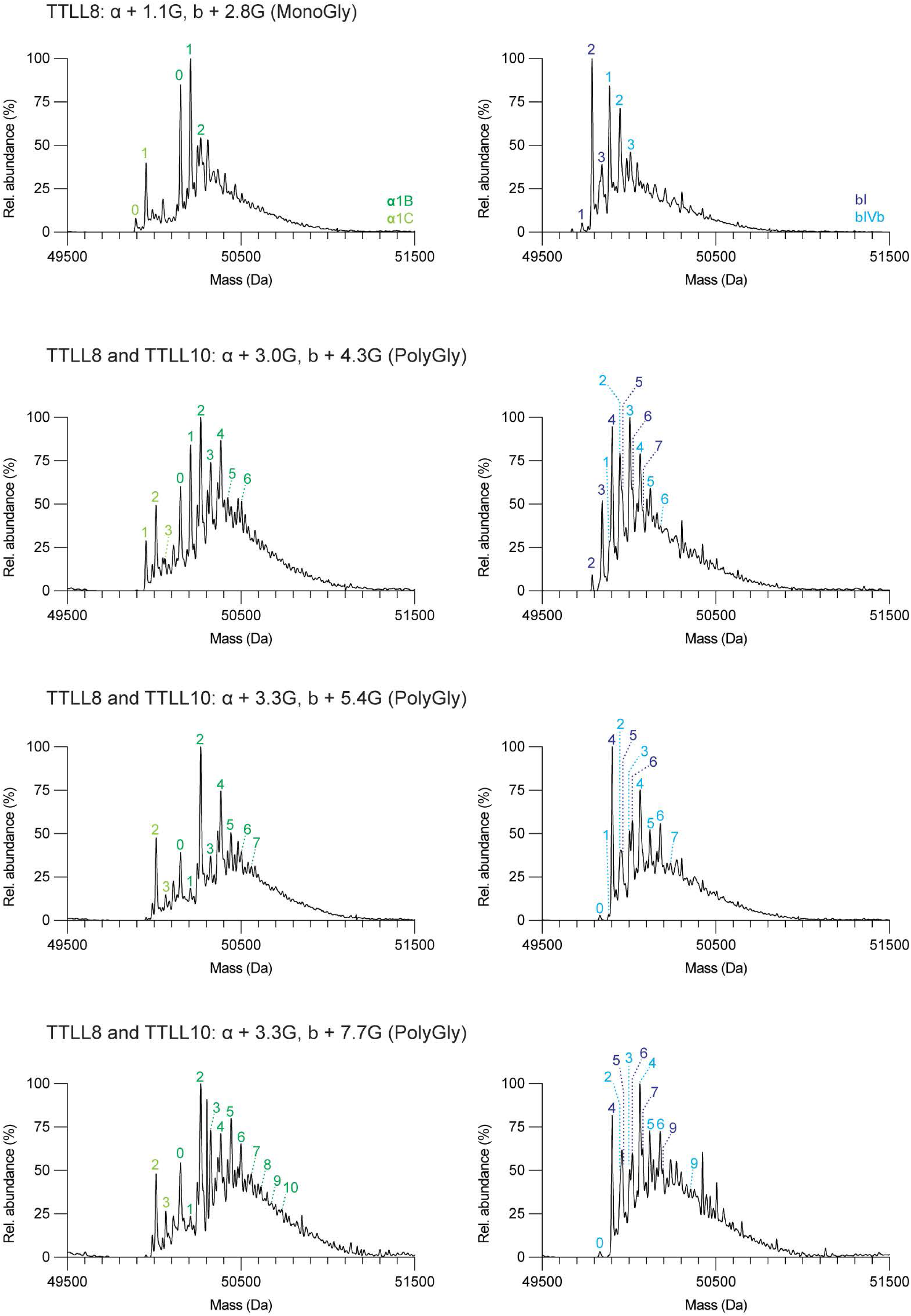
LC-MS of monoglycylated and polyglycylated microtubules used in binding assays shown in Figure 4. The weighted mean of the number of glycines (<n^G^>) added to α- and β-tubulin are denoted α + <n^G^>; β + <n^G^>.The number of posttranslationally added glycines is indicated in green (for α-tubulin isoforms) and blue (β-tubulin isoforms) on the spectra.

**Figure S8.**
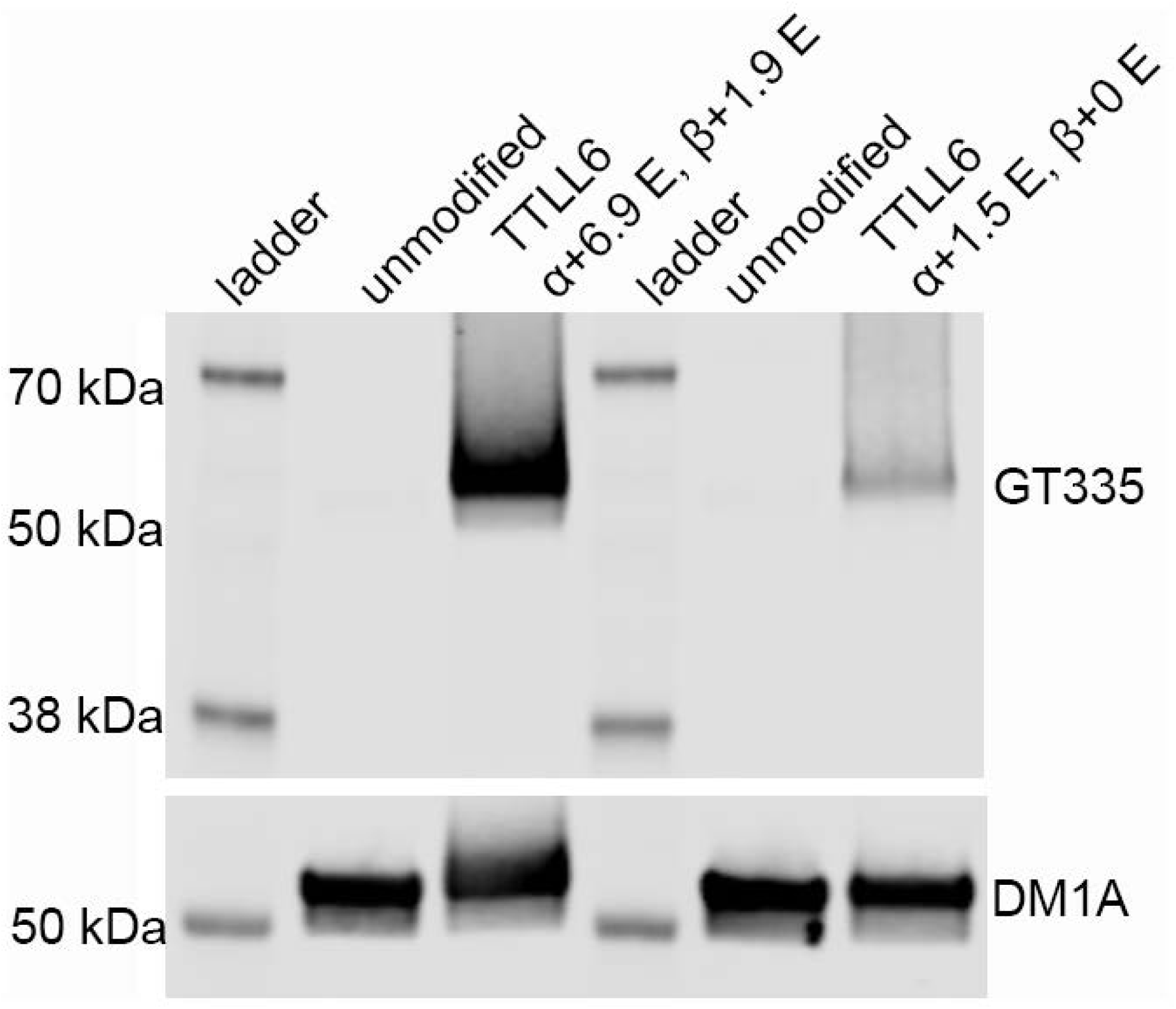
Western blot analysis of glutamylation in tubulin purified from tSA201 cells. 0.2 μg of tubulin was loaded for each unmodified and glutamylated sample. Glutamylation was detected with the GT335 antibody (Materials and Methods). No signal is detectable in the unmodified tsA201 purified tubulin. The MS/MS analysis of this tubulin did not identify any glutamylated peptides, either. In contrast, the tSA201 tubulin enzymatically glutamylated *in vitro* shows strong signal that increases with glutamylation level. Number of posttranslationally added glutamates indicated and determined from LC-MS measurements. The glutamylated tubulin runs slightly higher due to the changes in electrophoretic mobility.

**Figure S9.**
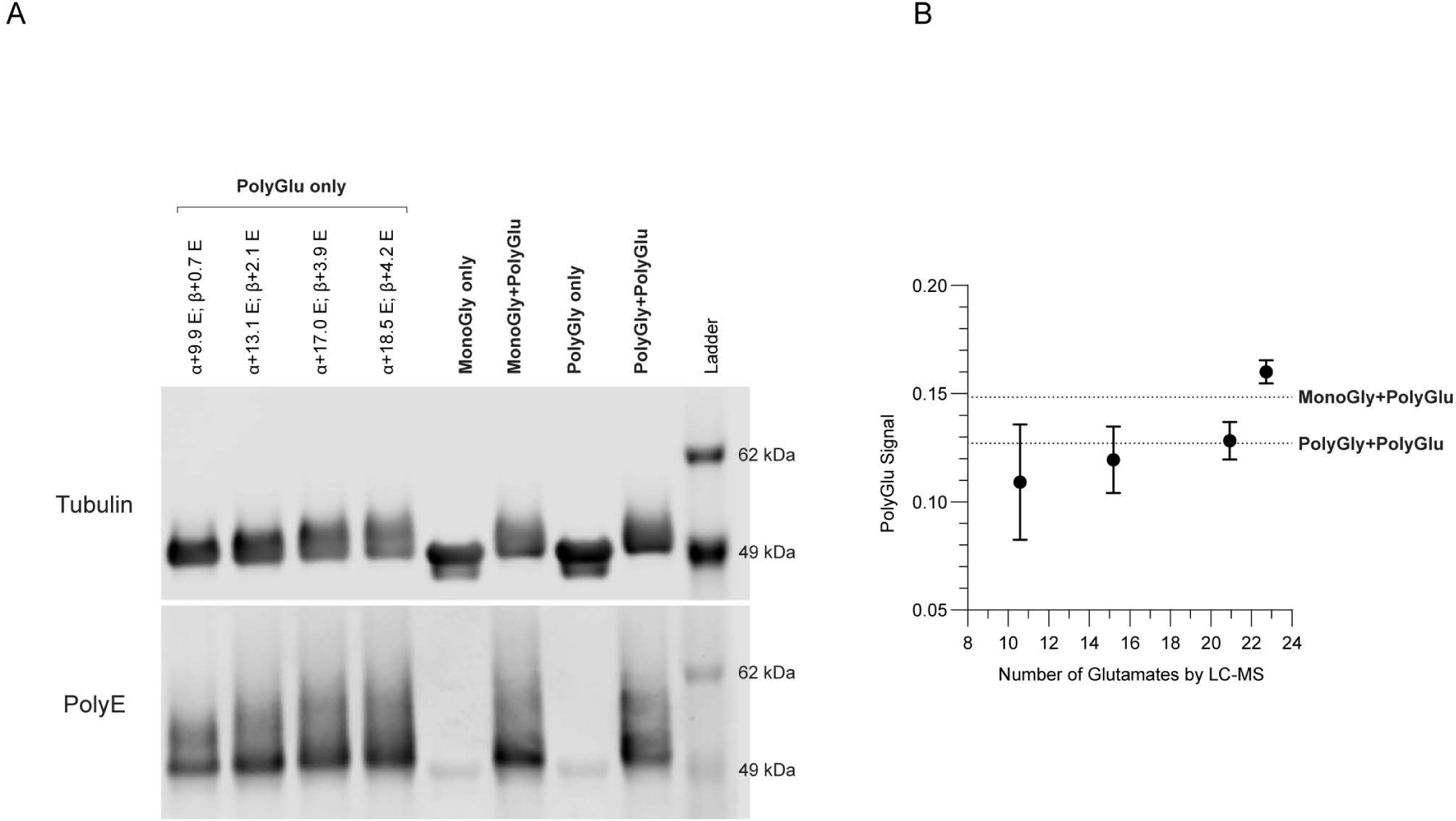
Estimation of glutamylation levels of monoglycylated and polyglycylated microtubules used in microtubule binding assays shown in Figure 5. (A) Western blot of polyglutamylated and monoglycylated or polyglycylated microtubules used in microtubule binding assays shown in Figure 5. From the left, first four lanes, TTLL6 polyglutamated microtubules with listed mean glutamate numbers determined from LC-MS spectra of intact microtubules. These were used to calibrate the polyglutamate signal detected with an anti-poly-E antibody (clone IN105, Adipogen; Materials and Methods); Subsequent four lanes, dually modified microtubules (glycylated with TTLL8 or TTLL8+10, and glutamylated with TTLL6) used in the assays shown in Figure 5; (B) Poly-E signal as a function of total glutamate numbers on α- and β-tubulin. The signal for the two dually modified species (monoglcylated + polyglutamylated and polyglycylated + polyglutamylated) used in the TIRF-based assays is shown with a discontinuous line.

## Materials and Methods

### Protein expression and purification

*Homo sapiens* TTLL8 (res. 39-585) was cloned into pCoofy28 (<u>Addgene plasmid #44004</u>) by sequence-specific ligation with RecA (44), and bacmid was produced by transformation of DH10EMBacY (Geneva Biotech). Baculovirus was produced by transfection of and subsequent amplification in ExpiSf9 cells (Thermo Fisher Scientific). TTLL8 was expressed in baculovirus-infected Sf9 cells (Thermo Fisher Scientific) with an N-terminal Glutathione S-transferase tag. Cells were infected at a density of 2.5×10^6^ cells/mL with an MOI of 2 and then grown for 48 hrs before harvest. Cells were lysed using a microfluidizer and cell lysate was clarified by centrifugation at 45,000 rpm (235,000 rcf) for 60 minutes using a Type 45 Ti rotor (Beckman-Coulter). The fusion protein was captured using GST affinity chromatography. TTLL8 was liberated from the resin using 3C protease at a 1:500 molar ratio, further purified on a heparin column, and finally polished by size-exclusion chromatography. TTLL8 was flash-frozen in small aliquots in 50mM HEPES pH 7.0, 10mM MgCl_2_, 200mM KCl, 2mM TCEP, 10% glycerol.

*Xenopus tropicalis* TTLL10 (res. 105-570) was cloned into pCoofy28 (<u>Addgene plasmid #44004</u>) and bacmid was produced by transformation of DH10EMBacY (Geneva Biotech). TTLL10 was expressed and purified following the same protocol as the one described above for TTLL8. TTLL10 was flash-frozen in 20mM Tris pH 8.0, 10mM MgCl_2_, 100mM NaCl, 2mM TCEP, 10% glycerol. TTLL10 with a C-terminal SNAP tag was expressed and purified as described above; however, the protein was conjugated to a SNAP-Surface Alexa Fluor 647 dye (New England Biosciences) after elution from the heparin column by incubation with the fluorophore on ice overnight. Excess dye was removed by size-exclusion chromatography whereupon TTLL10-SNAP(647) was flash-frozen in 20mM Tris pH 8.0, 10mM MgCl_2_, 100mM NaCl, 2mM TCEP, 10% Glycerol.

Unmodified human tubulin from tsA201 cells was purified as previously described using a TOG1 affinity column (27).

### Generation of glycylated microtubules for mass spectrometric analyses

The preparation of unmodified microtubules for this study closely follows previously published protocols (27, 30). Microtubules were polymerized from 10μM unmodified human tubulin in BRB80 (80mM PIPES, pH 6.8, 1mM MgCl_2_, 1mM EGTA) with 10% DMSO and 1mM GTP. Taxol was added to 20μM after 1hr, and then left to incubate overnight. Free tubulin was removed by spinning the polymerization mixture through a 60% glycerol cushion in BRB80 and the microtubules were resuspended in BRB80 with 20μM Taxol to a final concentration of 20μM. Glycylation reactions were performed at room temperature and commenced by addition of enzyme after the microtubules were evenly dispersed in reaction buffer. C^13^ glycine was used to improve separation of the modified βI and βIVb tubulin peaks by LC-MS and facilitate the identification of glycine-modified peptide fragments by MS/MS. After every glycylation reaction, bound glycylase enzyme was removed with a high salt wash by adding 300mM KCl, incubating at 37°C for 15 minutes, and subsequently spinning through a 60% glycerol cushion in BRB80 (80mM KOH-PIPES, pH 6.8, 1mM MgCl_2_, 1mM EGTA). For monoglycylation reactions, unmodified human microtubules stabilized with 20μM taxol were incubated with TTLL8 (0.2μM) at a 1:50 enzyme:tubulin molar ratio for 20 minutes at room temperature. For polyglycylation reactions with TTLL10, taxol-stabilized, unmodified human microtubules were first monoglycylated for 4 hours by TTLL8 (0.2μM) at a 1:20 enzyme:tubulin molar ratio, to ensure complete monoglycylation of the substrate. Bound TTLL8 was removed by a salt wash as described above, whereupon the monoglycylated microtubules were subsequently treated with TTLL10 (0.2μM) at a 1:50 enzyme:tubulin molar ratio for 20 minutes at room temperature. Bound TTLL10 was removed by a second salt wash performed as described above.

### Mass spectrometric analyses of glycylation reactions

The number of glycines added to α- and β-tubulin was determined by LC-MS. Reaction samples for LC-MS were prepared as described above and then quenched at 20 minutes *via* addition of 20mM EDTA on ice. Samples for LC-MS were diluted to 0.2μg/μL in cold BRB80 and then diluted 1:1 with 0.1% trifluoroacetic acid (TFA) in 20% acetonitrile. 1μg of microtubules were separated using a 0-70% acetonitrile gradient in 0.05% TFA at a flow rate of 0.2mL/min. The column was coupled to a 6224 ESI-TOF LC-MS (Agilent) and the spectra were deconvoluted using the Agilent MassHunter software and display a distribution of masses with consecutive glycylated species separated by 58 Da corresponding to one [^13^C]-glycine. The extent of tubulin glycylation on α- or β-tubulin was determined by calculating the weighted average of peak intensities for each tubulin species present.

Samples for MS/MS analyses were reduced with 5 mM Tris(2-carboxyethyl)phosphine (TCEP) hydrochloride at room temperature for 1hr, alkylated with 10 mM N-Ethylmaleimide for 10 min. Each sample was divided into 2 aliquots and digested with trypsin (Trypsin Gold, Mass Spectrometry Grade, Promega) and AspN (Sequencing Grade, Roche) separately. Trypsin and AspN digestion was performed with 1:15 enzyme:sample (w/w) at 37 °C for 12 hr. Digested samples were desalted using µElution HLB plate (Waters). Data acquisition was performed on a system where an Ultimate 3000 HPLC (Thermo Scientific) was coupled to an Orbitrap Lumos mass spectrometer (Thermo Scientific) via an Easy-Spray ion source (Thermo Scientific). For each LC-MS/MS run, 0.5 µg digests were injected. Peptides were separated on an ES902 Easy-Spray column (Thermo Scientific). The composition of mobile phases A and B was 0.1% formic acid in HPLC water, and 0.1% formic acid in HPLC acetonitrile, respectively. The mobile B amount was increased from 3% to 27% in 66 minutes at a flowrate of 300 nL/min. Thermo Scientific Orbitrap Lumos mass spectrometer was operated in data-dependent mode. The MS1 scans were performed in orbitrap with a resolution of 120K at 200 m/z and a mass range of 400-1500 m/z. Collision-induced dissociation (CID) method was used for MS2 fragmentation. MS2 scans were conducted in ion trap. The precursor ion isolation width was 1.6 m/z, and the dynamic exclusion window was 4 sec. Database search was performed using Mascot against Sprot Human database. The mass tolerances for precursor and fragment were set to 10 ppm and 0.6 Da, respectively. Up to three missed cleavages were allowed for data obtained from trypsin digestion, and up to four missed cleavages were allowed for AspN data. NEM on cysteines was set as fixed modification. Variable modifications include Oxidation (MW), Met-loss (Protein N-term), Acetyl (Protein N-term), 1hG(G), 2hG(G), 3hG(G), and 4hG(G). Peptides matched with heavy glycine modification were manually curated.

### Generation of differentially glycylated microtubules for microscopy assays

Unmodified human microtubules were polymerized from 10μM human unmodified tubulin with 1.5% biotinylated brain tubulin in BRB80 by addition of 10% DMSO and incubation for 1hr before adding 10μM taxol. These microtubules were modified with TTLL8 alone or with both TTLL8 and TTLL10. The mean number of glycines was determined by LC/MS. Specifically, to prepare differentially monoglycylated microtubules, 10μM umodified microtubules were incubated at room temperature with 1:20 TTLL8 for 1hr (<n^G^>_α_ ∼ 0.8, <n^G^>_β_ ∼ 1.2), 2hr (<n^G^>_α_ ∼ 1.1, <n^G^>_β_ ∼ 2.5), and 4hr (<n^G^>_α_ ∼ 1.1, <n^G^>_β_ ∼ 2.8). To prepare differentially polyglycylated microtubules, unmodified microtubules were first modified at room temperature for 4hrs with 1:20 TTLL8 (<n^G^>_α_ ∼ 1.1, <n^G^>_β_ ∼ 2.8). After removing TTLL8 using a salt wash as described above and subsequently spinning through a glycerol pad, these microtubules were treated with 1:20 TTLL10 for 2 (<n^G^>_α_ ∼ 3.0, <n^G^>_β_ ∼ 4.3), 5 (<n^G^>_α_ ∼ 3.3, <n^G^>_β_ ∼ 5.4), and 15 minutes (<n^G^>_α_ ∼ 3.3, <n^G^>_β_ ∼ 7.7). To prepare microtubules that were glycylated and glutamylated, unmodified microtubules were first modified with TTLL8 at room temperature for 1hr at a molar ratio of 1:50 (<n^G^>_α_ ∼ 0.1, <n^G^>_β_ ∼ 0.8) whereupon TTLL8 was removed by using a salt wash and spinning through a glycerol pad as described above. A portion of these microtubules were then modified at room temperature for 1hr with 1:50 TTLL10 (<n^G^>_α_ ∼ 0.4, <n^G^>_β_ ∼ 1.9) whereupon TTLL10 was removed by using a salt wash and spinning through a glycerol pad as described above. The monoglycylated microtubules and the polyglycylated microtubules were then split in half, with one half left untreated and the other half treated at room temperature overnight with 1:10 TTLL6 to obtain long polyglutmate chains primarily on α-tubulin (28), after which TTLL6 was removed through another salt wash and by spinning through a glycerol pad as described above. We note that we first glycylated and then glutamylated to ensure that we have the exact same glycylation levels for the unglutamylated and glutamylated microtubules. Glutamylation levels were estimated by western blot for glycylated/glutamylated microtubules, as it is not possible to quantify modification levels using LC-MS due to the large number of species present in the product mixture. Blots were stained first with a mixture of 1:10,000 rabbit anti-polyE antibody (clone IN105, Adipogen) and 1:10,000 mouse anti-α-tubulin antibody (clone DM1A, Sigma-Aldrich) for 1hr, washed, stained with a mixture of 1:10,000 IRDye ® 680RD goat anti-mouse antibody (LI-COR) and 1:10,000 IRDye ® 800CW goat anti-rabbit antibody (LI-COR), washed again, and then imaged on an Odyssey imager (LI-COR).

For Figures S3 and S8, microtubule samples were subjected to SDS-PAGE and Western Blot transfer. 200 ng of tubulin was loaded for each unmodified and glutamylated and glycylated sample. The membranes were blotted for modifications with the following antibodies: glutamylation with mouse GT335 antibody (AdipoGen Cat# AG-20B-0020-C100) at dilution 1:4000, mono-glycylation and bi-glycylation with rabbit Gly-pep1 (AdipoGen Cat# AG-25B-0034-C100) at dilution 1:2000, polyglycylation with the poly-Gly rabbit polyclonal antibody at dilution 1:4000 (45). Tubulin loading controls were detected with mouse anti-α-tubulin antibody (DM1A) (Acbam Cat# ab7219) at dilution 1:4000. Western Blots were visualized with secondary antibodies: IRDye^®^ 800CW Goat anti-Rabbit IgG Secondary Antibody (LiCor Cat# P/N: 926-32211) and IRDye^®^ 680RD Goat anti-Mouse IgG Secondary Antibody (LiCor Cat# P/N: 926-68070) at dilution 1:18000. All antibody dilutions were made in 4% milk in phosphate buffered saline with 0.2% tween.

### TIRF based microtubule binding assays

Flow chambers were constructed of plasma-cleaned and silanized glass slides and coverslips as previously described (46). Chambers were coated with neutravidin and then washed with BRB80 with 2mg/mL casein and 20μM taxol followed by BRB80 with 1% Pluronic F-127 and 20μM taxol. Microtubules were immobilized in the chambers and washed again with BRB80 with 2mg/mL casein and 10μM taxol followed by BRB80 with 1% Pluronic F-127 and 10μM taxol and finally equilibrated in binding buffer (60mM KCl, 10mM NaCl, 40mM PIPES pH 6.8, 2mM Tris pH 8.0, 20mM Glucose, 1.5mM MgCl_2_, 1.5mM TCEP, 1mM ATP 0.5mM EGTA, 1% Pluronic, 10μM Taxol). Taxol-stabilized human unmodified microtubules assembled with 1.5% biotinylated tubulin (Cytoskeleton, Inc. #T333P) and 1% HiLyte Fluor^TM^ 488-labeled tubulin (Cytoskeleton, Inc. #TL488M) were used as an internal control in all chambers for all assays. Microtubule binding assays were performed at 500nM TTLL10-SNAP(647) and were supplemented with oxygen scavenger mix to remove free oxygen from solution (7.5μM glucose oxidase, 0.5μM catalase). TTLL10-SNAP(647) was perfused into the chamber during image acquisition. Background-corrected line scan intensities were measured over time using Fiji and normalized to microtubule length and unmodified microtubules which were used as an internal standard in each chamber. Peak binding intensities were used for all microtubules.

Fluorescence images were acquired using an inverted total internal reflection fluorescence (TIRF) microscope (Nikon Ti-E with TIRF attachment) equipped with an iXON3-897 EMCCD camera (Andor). For TIRF assays, the excitation light was provided by a 488nm (Coherent Sapphire) or 647nm (Coherent Cube) laser set to 20mW before being coupled into the microscope *via* an optical fiber to the Ti-TIRF arm on the microscope. Light was delivered to the sample through the TIRF arm and directed towards a 100x 1.49 NA TIRF objective (Nikon CFI Apo TIRF 100x). Emission light was split using a dichroic beamsplitter FF640-FDi02 (Semrock) and further filtered using a FF01-550/88 filter (Semrock) on the 488 channel or a BLP01-647R filter on the 647nm channel. Prior to perfusing TTLL10, an image of all microtubules was taken using interference reflection microscopy (IRM) using an ORCA-Flash4.0 CMOS camera (Hammamatsu) and an image of the unmodified HiLyte 488-labeled, internal reference microtubules was taken by TIRF at 100ms exposure with excitation at 488nm. TTLL10-SNAP(647) binding during and after perfusion was imaged by TIRF with 100ms exposure at 2s intervals with excitation at 647nm.

